# T2T-CHM13 reference genome reduces mapping bias and enhances alignment accuracy at disease-associated variants

**DOI:** 10.64898/2025.12.17.694618

**Authors:** Ilaria Cherchi, Francesco Orlando, Orsetta Quaini, Marta Paoli, Yari Ciani, Francesca Demichelis

## Abstract

The T2T-CHM13v2.0 reference genome added previously uncharacterized genomic sequences and improved the accuracy of repetitive stretches compared to former human genome assemblies. By comprehensive allelic variation analysis and read mapping statistics from sequencing reads aligned to hg38 and T2T-CHM13 assemblies in samples encompassing different sequencing designs and ethnicity groups, we observed that T2T-CHM13v2.0 assembly significantly reduces the reference mapping bias (RMB) and increases read mapping precision at clinically relevant sites, including *BRCA1* pathogenic variants. Further, we report the presence of sequence dissimilarities among reference genomes in the proximity of ClinVar annotated variants, suggesting the need for data re-analysis and potential redesign of probes targeting clinically relevant regions. Overall, these findings support the implementation of T2T-CHM13 reference for the improvement of sequencing data analyses in the clinical genomic setting.

## 2 Introduction

The first telomere-to-telomere (T2T) reference genome sequenced from the CHM13 human cell line uncovered around 200 million base pairs of genomic sequence, including the short arms of acrocentric chromosomes and centromeric satellite arrays^1^. The T2T-CHM13v1.0 assembly corrected falsely duplicated hg38 reference genome regions, leading to improved variant calling across multiple populations^2^. Additionally, the v2.0 version provided chromosome Y integration^3^ and expanded the study of repeat elements^4^, epigenetic features^5^, and structural variants^6^, revealing large-scale differences with previous human genome assemblies^7^. Since variations in the nucleotide sequence of a reference genome can considerably affect sequencing read mapping, we sought to specifically assess the effect of aligning reads to T2T-CHM13 - instead of to hg38 - for performing variant allele frequency (VAF) estimation at single nucleotide polymorphisms (SNPs) and for the analysis of clinically relevant variants, CpG islands, and transcription factor binding sites (TFBS).

Further, we reasoned that T2T-CHM13 could significantly ameliorate the bias towards higher mapping rates of the reads supporting the reference allele with respect to the alternative allele at heterozygous SNPs, defined as read-mapping bias or reference mapping bias (RMB)^8^. RMB affects the estimation of allelic frequency at hypervariable sites, e.g., HLA genes^9^, and annotated variants, preventing a precise quantification of either inherited or acquired allelic imbalance. Methods developed for mitigating RMB include the alignment of sequencing reads to a global major-allele reference^10^ or pangenome^11^, and the correction of VAF estimates^12^.

The inflation of reads supporting the reference allele observed in RMB has been attributed to mapping errors caused by the variability among the selected reference genome and the individual’s genomic sequence in the context of ancient DNA analysis^13^. We hence hypothesized that the alignment of human reads to the T2T-CHM13 reference genome would ameliorate the estimation of heterozygous SNPs VAF without requiring RMB correction or additional pre-processing steps.

To evaluate the impact of the T2T-CHM13 assembly on the alignment of reads and on the presence of reference mapping bias, we performed in-depth characterization of sequencing data from multiple sequencing designs. We showed that the alignment to T2T-CHM13 reference significantly enhances allelic variation estimation and read mapping when compared to previous assemblies (e.g., hg38). Mismatches detected in the genomic sequences in the proximity (±150bp) of SNPs with discordant VAFs across the two assemblies demonstrate that RMB is caused by genomic inaccuracies in the reference genome. We performed an annotation of sequence dissimilarities across reference genomes in UCSC CpG islands, TFBS and ClinVar variants proximities’ for a prompt evaluation of the necessity for data re-analysis and potential re-design of sequencing probes for clinical applications. We envision that T2T-CHM13 assembly read alignment will facilitate the detection of low VAF variants relevant to precision medicine testing via increased VAF and local coverage at disease-associated genomic sites, possibly improving variants detection.

## 3 Results

### 3.1 Reference mapping bias (RMB) is significantly reduced in samples aligned to T2T-CHM13

For testing whether aligning reads to the T2T-CHM13 reference genome would reduce RMB and, consequently, affect VAF computation, we aligned a total of 89 samples to hg38 and T2T-CHM13v2.0 assembly and generated pileups on a genome-wide set of 30 million high minor allele frequency (MAF) SNPs (**Figure 1A**). To provide a comprehensive overview of multiple state of the art sequencing designs, the study included: 25 representative genomic DNA (gDNA) whole-genome sequencing (WGS) data from the 1000 Genomes Project (1kGP; median coverage 30x)^14^, 8 tumor cell line gDNA data from the Cancer Cell Line Encyclopedia (CCLE) WGS^15^ (median coverage 30x), 26 matched gDNA and cell-free DNA (cfDNA) samples from breast cancer patients sequenced with whole-exome sequencing (WES; median coverage 96x for gDNA and 500x for cfDNA)^16^, and 30 gDNA and cfDNA samples from prostate cancer patients profiled with a prostate-cancer specific targeted assay (PCF_SELECT^12^; intended coverage of 200X and 1000X for gDNA and cfDNA, respectively). Additionally, hg38 and T2T-CHM13 alignment files from N=1017 individuals from the recent Oxford Nanopore Technologies (ONT) sequencing of 1kGP^17^ were analyzed to provide an overview on the impact of reference genome change in long-read sequencing data. First, comparative analysis of read alignment metrics was performed on a sample basis for hg38- and T2T-CHM13 aligned data (**Supplementary Figure 1A**). Previously reported differences in mapping statistics for T2T-CHM13 versus hg38 aligned samples^2,18^ are overall in line with observed trends among all sequencing designs (**Figure 1B**). In particular, a marked increment in read average quality (mean of 5% increase), and a decrease in error rate (mean of 10.7% decrease) was registered (**Figure 1B**).

**Figure 1.**
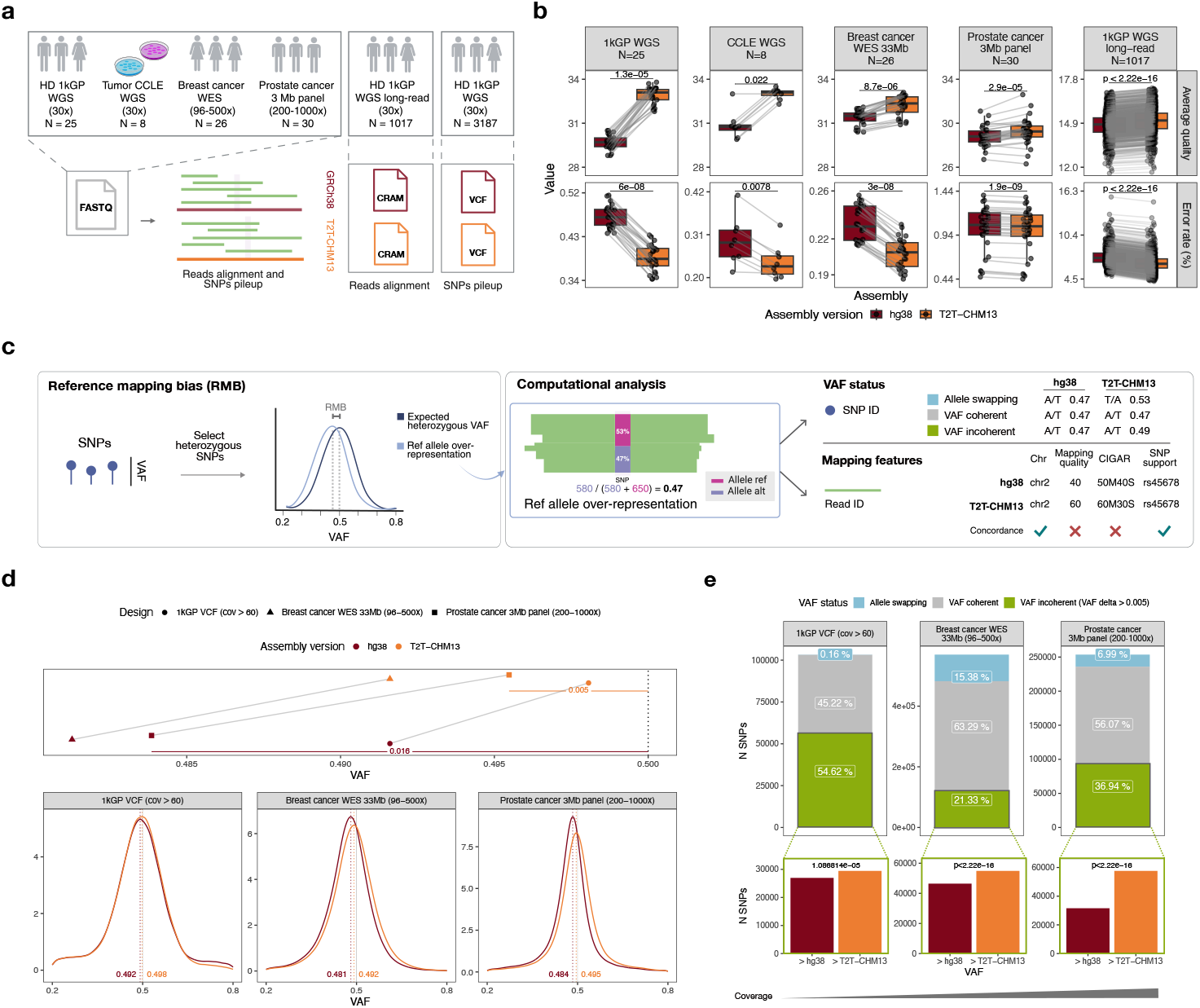
Alignment to T2T-CHM13v2.0 reference genome improves mapping metrics and significantly reduces RMB across diverse sequencing designs. a) Sketch summarizing the cohorts analyzed in this study. b) Paired mapping statistics for all hg38 and T2T-aligned samples. Each line connecting dots (jitter applied for visualization purposes) corresponds to a sample. Wilcoxon paired test significance is shown for each design/metric pair. c) Sketch of RMB (left) and computational analysis performed to explore VAF status changes and mapping features of reads spanning SNPs (right). [Created with BioRender] d) Reference mapping bias analysis for 1kGP VCF entries shared among hg38 and T2T-CHM13 with at least 60 supporting reads, WES and panel-based sequencing samples. The top panel compares peak values for hg38 and T2T with respect to the expected threshold of 0.5, highlighted by a dotted vertical line. Difference between the expected threshold and median peak value among cohorts is reported for both hg38 and T2T-CHM13 alignment. The bottom panel includes the densities of all analysed heterozygous SNPs VAF values and reports the numeric value of each density peak, corresponding to the dotted vertical lines. The expected VAF value of 0.5 is indicated by the continuous gray vertical line and the value for the RMB peak is depicted by dotted lines colored by assembly. e) Barplots provide a classification for each heterozygous SNP based on VAF comparison among hg38 and T2T-CHM13. The focus on SNPs with incoherent VAF among assemblies reports the number of entries with VAF increase for hg38 (in dark red) and T2T-CHM13 (orange). One-tailed Wilcoxon test for VAF T2T-CHM13 > VAF hg38 are reported on the top of the bars. The annotation on the bottom highlights increasing coverage among the cohorts.

The extent of RMB as a function of the reference genome considered was assessed by comparing the VAF distributions of heterozygous SNPs with respect to the expected threshold at 0.5, and annotating SNP features among hg38- and T2T-aligned samples, including VAF status across assemblies and comparison of SNP supporting reads (**Figure 1C**). While comparing intra-individual VAF standard deviation and coverage over high minor allele frequency (MAF) heterozygous SNPs (i.e., MAF > 0.2 across all major ethnicities and 0.2 ≤ VAF ≤ 0.8; **Supplementary Table 2**), larger VAF differences were observed at SNPs with lower coverage values (**Supplementary Figure 2A**). In concordance with trends observed both in theoretical and observed VAF distributions at varying coverage thresholds (**Supplementary Figure 2B**), samples sequenced at 30x coverage (i.e., WGS 1kGP and CCLE samples) did not exhibit RMB (**Supplementary Figure 2C**), albeit still underlying the presence of several SNPs with varying VAF among the two assemblies (18.5%). To expand the study of RMB to higher coverage data from WGS samples, we extracted VAF estimations from the 1kGP VCFs genotype calls (n=3202 individuals) for hg38^14^ and T2T-CHM13v2.0^2^. We retained only positions annotated as high MAF SNPs, present in both VCFs, and supported by more than 60 reads (i.e., local coverage > 60). VAF distributions from 1kGP VCFs showed a peak closer to the expected heterozygous value in T2T-CHM13 estimates compared to hg38 (0.498 and 0.492, respectively – bootstrap p-value = 2e-04) (**Figure 1D**). Focusing on high-coverage samples from WES and custom targeted panel (∼3Mbp) sequencing, we observed that heterozygous SNPs VAF density curves from samples aligned to T2T-CHM13 were centered around the expected 0.5 value, *de facto* abolishing RMB. In contrast, hg38-aligned samples exhibited a peak at 0.48, indicating a preference for the reference allele that is concordant with the presence of RMB.

SNPs VAF status was classified as incoherent VAF and coherent VAF when the absolute VAF difference (delta VAF, i.e., |*VAF T*2*T* − *VAF hg*38|) among assemblies was greater and lower than 0.005, respectively. 54.62% of SNPs from 1kGP VCFs, 21.33% of SNPs in breast cancer WES samples, and 36.94% of SNPs in prostate cancer samples sequenced with the custom targeted panel were classified as sites with incoherent VAF (**Figure 1E**), underlining the high extent of variation among estimated VAF values from hg38 to T2T-aligned data. A global significant increase in the support towards the alternative allele for T2T-CHM13 was observed at sites with incoherent VAF among assemblies (p-value = 2.808e-09 for 1kGP VCF data and p-value<2.22e-16 for WES and custom targeted panel data one-tailed Wilcoxon paired test for VAF T2T-CHM13 > VAF hg38), with a higher number of SNPs with VAF increase in T2T-CHM13 alignment observed in samples with higher coverage values (**Supplementary Figure 2D**). RMB shifts and differences in VAF status were also observed when stratifying 1kGP VCF data by super-population groups (**Supplementary Figure 2E**).

To pinpoint the cause of the improved mapping on alternative alleles in T2T-CHM13, we analyzed the mapping features of the reads spanning heterozygous SNPs with consistent VAF status, i.e., incoherent VAF or coherent VAF classification in all breast cancer samples (**Supplementary Table 3**). Of the hg38 SNPs with consistent VAF status, 5.1% were either mapped to a different chromosome or resulted unmapped, and 18.8% were mapped to another position within the same chromosome when aligned to T2T (**Supplementary Figure 3A-B**). To determine if VAF shifts are also associated with mapping differences related to reference sequence variation, we extracted the DNA flanking sequences (± 150 bp) from each previously characterized SNP with a consistent VAF status. We performed pairwise alignment among T2T-CHM13 and hg38 sequences (**Methods 5.8**). SNPs supported by at least one different sequencing read (discordant support) exhibited a significantly higher proportion of sequence mismatches with respect to SNPs with concordant support (Wilcoxon one-tailed test p-value = 2.4e-05, **Supplementary Figure 3D**). Overall, these data confirm that the reference genome sequence discrepancies result in differential mapping, whereby T2T-CHM13 globally reducing the difference between expected versus observed VAFs, thus abolishing the RMB.

### 3.2 Comparison of genomic sequences from functionally annotated regions reveals differences between hg38 and T2T-CHM13

Since differences in sequences proximal to SNPs were linked to inconsistencies in mapping, we inquired whether other genomic annotated regions might suffer from mapping biases. We analyzed the UCSC tracks for CpG islands^19^, ENCODE ChIP TFBS peaks^20^ and the proximal sequence of variants annotated in the ClinVar^21^ database (± 150bp) through pairwise sequence alignment to annotate regions where sequence discrepancies occur among hg38 and T2T-CHM13 (**Figure 2A, Supplementary Table 4**). Observed mismatch rates range from 16.57% for flanking regions of ClinVar variants to 36.89% for CpG islands, while insertions and deletions (i.e., InDels) were present at a similar extent throughout the tracks (min: 1.89%, max: 6.45%; **Supplementary Figure 4A**). The same trends were observed when comparing hg19 to the T2T reference genome, while no considerable differences were observed between hg19 and hg38 (∼ 1% mismatches and < 0.5% insertions and deletions) (**Supplementary Figure 4A**).

**Figure 2.**
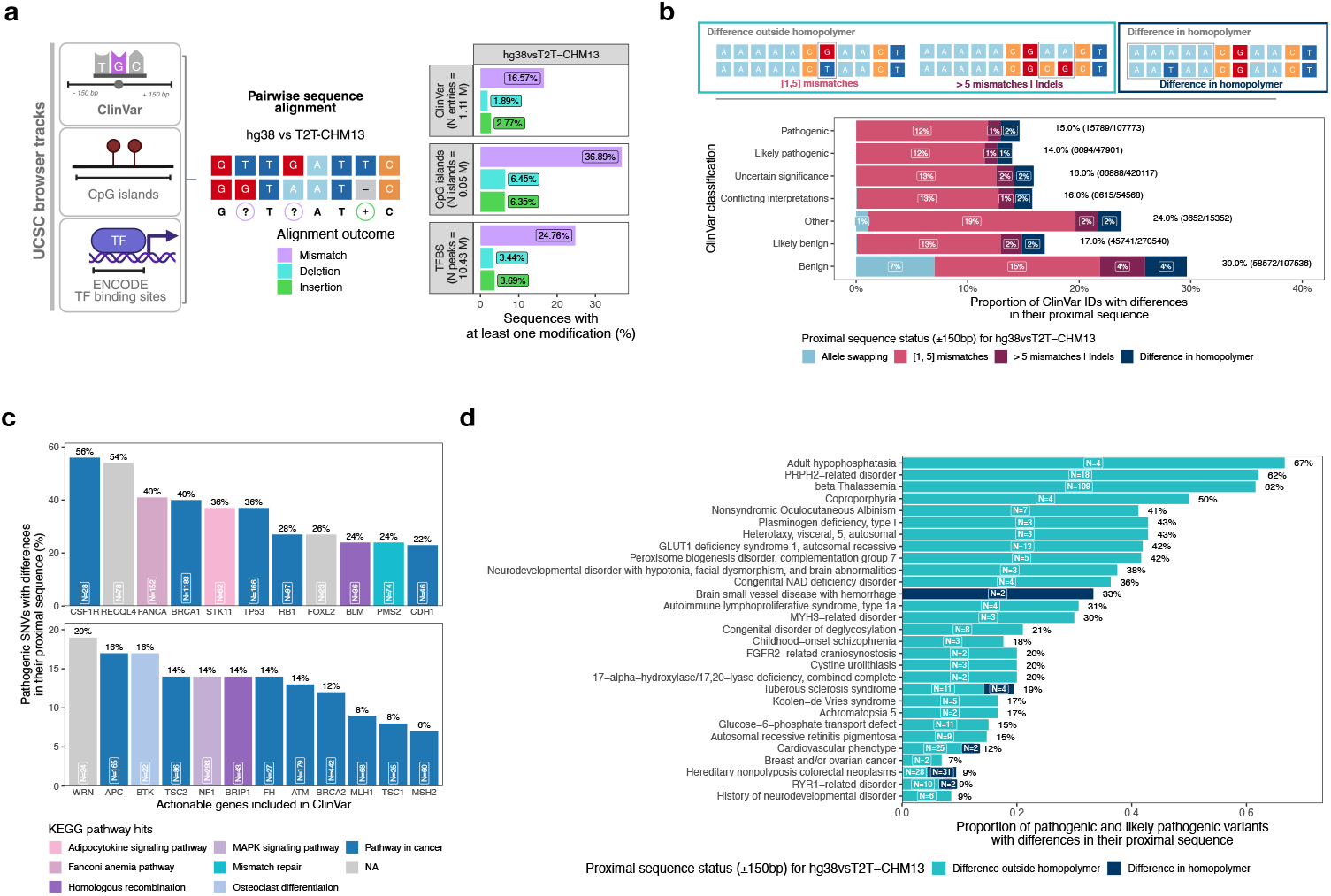
Comparison of genomic sequences from annotated regions reveals variation among hg38 and T2T-CHM13. a) Summary of pairwise alignment analysis of regulatory and genomic variation elements. Left: sketch presenting the genomic annotation tracks analyzed through pairwise alignment. [Created with BioRender] Right: barplot reporting the proportion of annotation tracks entries with at least one modification (insertions, deletions and mismatches) comparing hg38 and T2T-CHM13 sequences. b) Analysis of ClinVar variant’s sequence proximity for hg38vsT2T-CHM13 comparison. Top: sketch of sequence differences within and outside homopolymer stretches. Bottom: ClinVar IDs stratification based on pairwise sequence alignment classification and variant classification. The total percentage of entries with sequence differences is reported on the right of each barplot. c) Barplot reporting actionable genes included in COSMIC Actionability signature with at least 20% of ClinVar pathogenic variants with sequence differences in their proximity. The total number of variants with differences per gene is reported within each bar and the total percentage of pathogenic variants on the top of the bar. Bars are colored according to the top KEGG pathway hit per gene. d) Proportion of pathogenic variants by hg38vsT2T-CHM13 pairwise alignment analysis outcome for all MEDGEN disease type present in ClinVar with > 10 pathogenic variants and at least one variant with sequence differences.

Focusing on CpG islands, we quantified that 40.3% of CpGs have at least one base difference (either a mismatch or an InDel) between hg38 and T2T-CHM13 of which ∼ 63% show a different number of C bases (**Supplementary Figure 4B**). When assessing the potential impact on methylation analyses, we observed that 0.2% of the reads reported a change in methylation levels, either being cytosines or changes in the context (CpG, CHG, CHH), suggesting that differences in the assembly do not significantly impact downstream analyses (**Supplementary Figure 4C, Methods 5.10**). Negligible effects were observed when performing TF enrichment analysis of TF ChIP-seq peaks’ sequences (**Supplementary Figure 4D-4E, Methods 5.11**).

To investigate whether discrepancies in the proximal sequence of ClinVar variants were enriched within repetitive regions, we annotated the presence of homopolymeric repeats - defined as repetitive stretches (≥5bp) of the same base. Cumulatively, 81.8% of ClinVar variants’ proximal sequences (± 150 bp) harbor at least one homopolymer stretch, which varies among hg38 and T2T-CHM in 2.6% of variants (**Supplementary Figure 5A**), corresponding to 2.1% of all ClinVar variants. The analysis of the proximal DNA sequence of ClinVar annotated variants highlighted the presence of different nucleotide compositions among reference genomes for ∼17% of the variants (**Figure 2A**), with 15% of ClinVar pathogenic variants with at least one mismatch or indel in their proximity when comparing hg38 and T2T-CHM13 sequences (**Figure 2B**). In particular, 8% (24 out of 304) of the actionable genes included in the COSMIC Actionability v14^22^ list harbor more than 20 pathogenic variants with sequence discrepancy among hg38 and T2T-CHM13 in their proximity, suggesting that correct variant detection might depend on the choice of the reference genome. Among genes especially affected by these differences (i.e., > 5% of pathogenic variants with proximal sequence modifications and encompassing > 100 ClinVar entries), we identified genes involved in cancer pathways, e.g., *BRCA1, ATM, MSH2, CHEK2, FANCA, BRCA2*, as well as other disease-related pathways (**Figure 2C, Supplementary Figure 5C**).

We analyzed the MEDGENE code annotation within the ClinVar VCF file to extract disease information and observe the incidence of variation within and outside homopolymers among all pathogenic and likely pathogenic variants per disease (**Figure 2D**). Twenty-two annotated diseases, including PRPH2-disorder and beta thalassemia, have at least 15% of pathogenic and likely pathogenic variants with differences outside homopolymers.

### 3.3 Reference genome sequence differences within homopolymeric stretches help elucidate differences in SNV VAF estimates among hg38- and T2T-aligned samples

To evaluate the effect of the reference genome on the detection of clinically relevant SNVs, we performed variant calling on a set of cfDNA samples from prostate and breast cancer patients, using Strelka2^23^ and BCFtools^24^ (**Figure 3A, Supplementary Table 5**). Focusing on ClinVar sites with variations in their proximal sequence, both Strelka2 and BCFtools identified more pathogenic variants in patients’ data aligned to T2T-CHM13 with respect to hg38 (+ 22.6% and + 24.7% for Strelka2 and BCFtools, respectively; **Figure 3B**). Additionally, a significant increase in the quality of SNV calls for T2T-aligned samples (p-value < 2.22e-16) was observed for both variant calling methods. The consensus among variant callers over pathogenic and likely pathogenic variants was higher in T2T-aligned samples when compared to hg38-aligned with 508 (10%) and 484 (9.4%) shared calls among variant callers, respectively (**Supplementary Figure 6A**).

**Figure 3.**
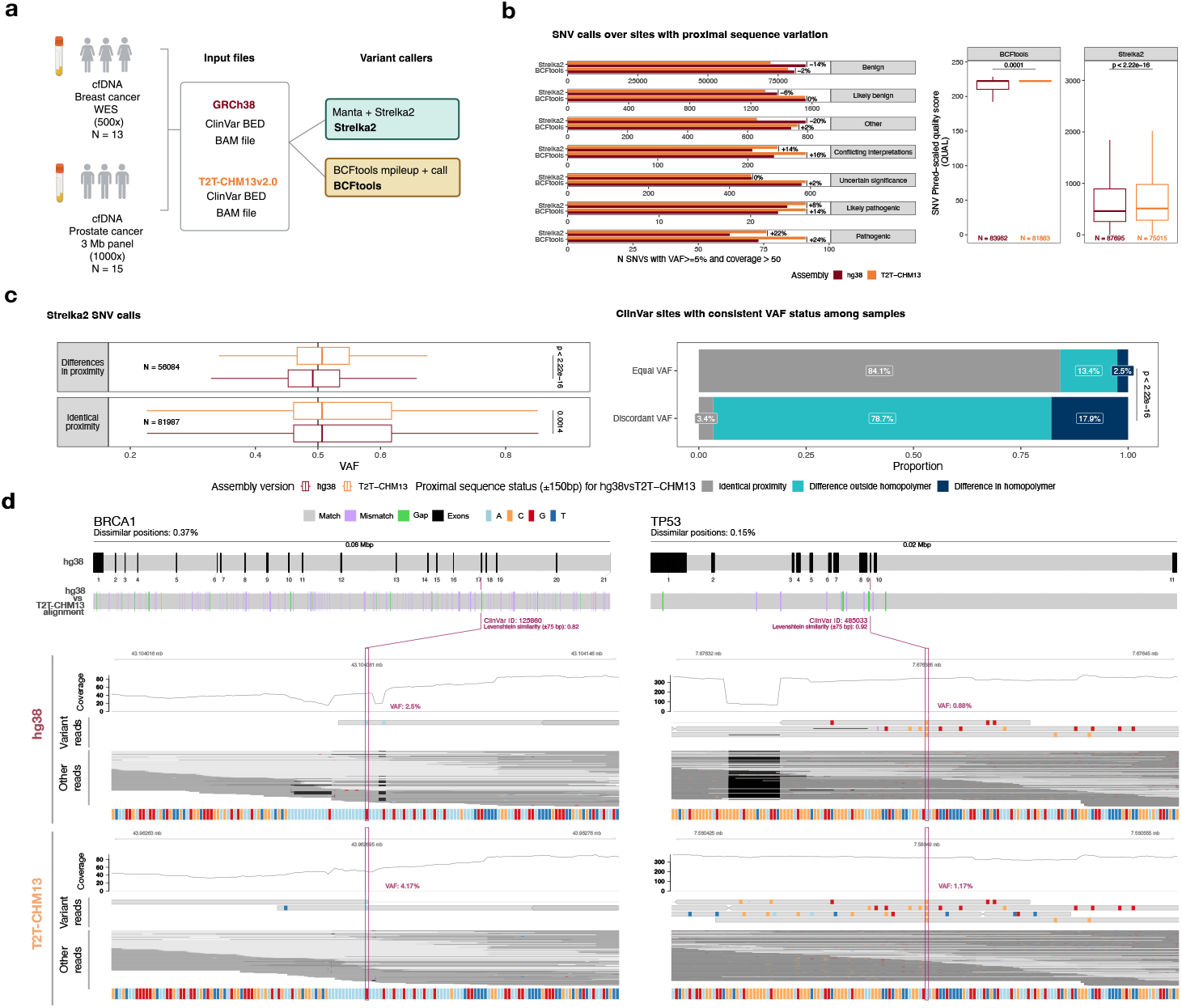
Variant calling over ClinVar sites shows improved SNV detection for samples aligned to T2T-CHM13. a) Schematic representation of variant calling analyses performed on cell-free DNA samples from tumor patients (WES and 3Mb panel) on ClinVar sites. [created with BioRender] b) SNV calls with at least 5% VAF and 50 supporting reads performed with Strelka2 and BCFtools. Left: number of identified SNVs stratified by ClinVar classification. Right: quality score for SNV calls performed with Strelka2 and BCFtools. One-tailed pairwise Wilcoxon test for T2T-CHM13 > hg38 p-value is reported for both variant calling methods. Outliers not shown. The number of data points is indicated at the bottom of each boxplot. c) Strelka2 variant calling. Left: boxplot for VAF estimation from variants with and without sequence differences in their 150 bp proximity. Wilcoxon test p-value was computed for all comparisons. Right: barplot annotation reporting the percentage of SNVs with differences within their homopolymeric stretch. Homopolymers were defined as stretches of ≥ 5 equal bases. d) IGV-like representation of differential read support over selected ClinVar variants in *BRCA1* (left) and *TP53* (right) gene. Top: gene exon structure (hg38 coordinates) and hg38vs T2T-CHM13 BLAST comparison of the entire gene sequence, colored by detected sequence differences. Total length of the analysed sequence and percentage of dissimilar bases are reported on the top. Bottom: genomic coordinates, coverage profile, reads supporting and not supporting ClinVar variant allele (Other reads), and reference sequence composition for hg38 and T2T-CHM13 reference genome. The variant of interest is highlighted in red and VAF estimate is reported for each alignment.

Importantly, we observed a significant increase in VAF estimates for T2T-aligned samples data when compared to hg38-aligned data at sites with sequence variation (one-tailed Wilcoxon test, p-value <2.22e-16), while entries with no modifications in their proximal sequence showed coherent VAF (**Figure 3C** and **Supplementary Figure 6B**). Analyzed variants not only include somatic SNVs but also inherited germline SNVs and can also coincide with annotated dbSNP sites, suggesting that the VAF improvement observed in samples aligned T2T-CHM13 aligned partly reflects RMB reduction. To remove potential sample-specific biases, we retained only SNVs with a consistent impact of reference genome change on VAF (i.e., VAF always coherent or incoherent between assemblies across all samples), and we observed a clear enrichment (Fisher test, p-value <2.22e-16) for sequence differences within and outside homopolymeric stretches across sites with discordant VAF (**Figure 3C**).

In-depth characterization of read alignment at ClinVar sites across regions with homopolymer variation highlighted a differential read support for hg38- and T2T-aligned samples, impacting VAF estimates and coverage homogeneity across well-characterized genes, as *BRCA1* and *TP53* (**Figure 3D**). Reads overlapping sites with diverging sequence among assemblies were classified as InDels upon hg38 alignment but not upon T2T-CHM13 alignment, suggesting that the sequence present in T2T-CHM13 assembly better recapitulates the correct sequence (i.e., present in the patient’s genome). The *BRCA1* gene structure is impacted by numerous changes between hg38 and T2T, which result in a modified read mapping both at exonic and intronic regions (**Supplementary Figure 6C**). Coverage increment was observed at likely pathogenic and pathogenic variants within *BRCA1* exon 17 for T2T-aligned samples.

Overall, 24% of genes reporting pathogenic or likely pathogenic variants in ClinVar exhibited increments in the VAF of their pathogenic or likely pathogenic variants located on regions of genomic sequence discrepancy (**Supplementary Figure 6D**), supporting the notion that adoption of the T2T-CHM13 reference genome improves VAF estimation at numerous genomic sites of clinical interest. To investigate whether the genomic sequence from diverse individuals at these sites is closer to T2T-CHM13 or hg38 reference sequence, we performed a read consensus analysis across 1,017 1kGP long-read individual sample data. Sequence similarity computed between read consensus sequences from the 1kGP long-read cohort and the reference genome over pathogenic or likely pathogenic variants was significantly higher for T2T-aligned samples with respect to hg38-aligned samples (One-tailed Wilcoxon paired test for T2T similarity > hg38 similarity p-value < 2.22e-16), further underlining the higher fidelity of T2T-CHM13 to individuals’ genomic sequences against hg38 at these sites irrespectively of super-population groups (**Supplementary Figure 6E**).

## 4 Discussion

The choice of the reference genome is crucial for analyzing sequencing data, as it significantly impacts read alignment accuracy and the outcome of downstream analyses. The T2T-CHM13 genome build enables significant advancements in genetics and genomics studies relevant to human diseases by uncovering previously uncharacterized regions through cutting-edge third-generation sequencing technologies and novel assembly methods. We systematically investigated whether significant differences would be observed in the analysis of genomic features when changing the reference genome.

Based on the analysis of both in-house and publicly available DNA sequencing datasets across diverse sequencing designs, we underlined the value of aligning human sequencing data to T2T-CHM13v2.0 reference to reduce RMB and globally enhance read mapping. Discrepancies in VAF estimates were consistently present across all analyzed samples and assay designs when mapped on hg38 in respect to T2T-CHM13: globally, aligning to the T2T-CHM13 reference genome improves read mapping and SNP VAF estimation, regardless of the sequencing strategy or intended coverage. We envision that the current efforts for building a comprehensive human T2T pangenome^25^ will further improve read mapping accuracy and aid in the characterization of the human genome landscape. Furthermore, the analysis of increasingly growing long-read sequencing datasets requires the alignment to complete human genomes for the identification of structural variants and complex genetic variations^17,26^.

Through pairwise sequence comparison analysis, we annotated sequence differences in CpG islands, TFBS peaks, and ClinVar sites, and we provide a useful resource for re-evaluating sites likely prone to differential mapping (available on Zenodo and as a ClinVar OpenCRAVAT^27^ annotation). In this study, while genomic differences within ChIP-seq peaks and CpG islands had no significant impact on TFBS enrichment analyses and methylation calling outcomes, coverage and VAF increment at ClinVar variants suggest that the latest reference genome could, in principle, provide an advantage for variant identification with respect to hg38. Indeed, realigning reads to T2T-CHM13v2.0 assembly might ameliorate the detection of well-characterized pathogenic variants included in genetic tests for therapy selection, particularly for variants with low VAF. As reported in a recent publication on variant detection from Swedish population sequencing data^18^, T2T-CHM13 alignment enhances the identification of disease-causing variations and improves read alignment metrics. In our study, by analyzing the sequences proximal to ClinVar annotated variants, we identified the presence of widespread discrepancies among reference genomes as the cause of the previously observed improvements in mapping accuracy at clinically relevant sites. In addition, sequence differences detected in homopolymer stretches and low-complexity regions among hg38 and T2T-CHM13 underscore the importance of reference genome choice for designing DNA probes, which could greatly benefit from a more accurate sequence for improving affinity and sequencing yield. Considering our observations, we encourage the adoption of T2T-CHM13 reference for human genomic data alignment, especially for the evaluation of clinically relevant variants.

### 4.1 Limitations of the study

This study assessed the impact of T2T-CHM13 assembly on individual features relevant to variant calling and precision medicine, both in the setting of targeted and whole genome sequencing. The lower median coverage of analyzed WGS samples (30x) affected the ability to identify peak shifts in VAF distributions, thus impeding a careful assessment of RMB in the genome-wide setting. Additionally, we expect that RMB analysis of long-read data from Oxford Nanopore Technologies and PacBio sequencing might yield different results with respect to trends observed in short-read data^28^. Further limitations include the necessity to rely on liftOver^29^ conversion for comparing the same sets of genomic regions among assemblies, which did not fully account for intra-reference diversity and prevented the exploration of novel sequences added into the T2T-CHM13v2.0 assembly. Finally, potential improvements in the limit of detection resulting from using T2T-CHM13 reference instead of hg38 should be evaluated in a more extensive set of samples and orthogonal validations.

## 5 Methods

### 5.1 Sample collection and sequencing

#### Breast cancer WES (96-500x)

Plasma samples were obtained from a cohort of breast cancer patients enrolled on a protocol approved by the Ethics Committee of Santa Chiara Hospital in Trento (Rep.Int.12315 of July 24, 2017) with written informed consent. Eligibility criteria included breast cancer diagnosis and recommendations for neoadjuvant therapies. Blood collection and subsequent cfDNA and gDNA extraction were performed as described in a previous publication^16^. For each patient, 20-50 ng of cfDNA and 100 ng of matched germline DNA (gDNA) sonicated to 180-220bp (Covaris M220) were utilized for DNA whole-exome sequencing library preparation and hybridization capture (KAPA HyperPrep Kit, SeqCap EZ HyperCap v. 2.3 Protocol; Roche). Sequencing was performed at the University of Trento by the Next Generation Sequencing Facility on Illumina HiSeq2500 platform with a paired-end protocol (100bp) with a mean coverage of 537x (282x - 731x) for cfDNA and of 96x for gDNA (89x - 109x).

#### Prostate cancer 3Mb panel (200-1000x)

Prostate cancer plasma and buffy coat samples were obtained from patients with prostate adenocarcinoma receiving androgen deprivation therapy from PRIME Observational Trial who provided informed consent. The PRIME Observational Trial (ClinicalTrials.gov ID: NCT06981377) received ethics approval by Santa Chiara Hospital in Trento on 31/01/2019 (Rep. Int. # 2562) and subsequently by the Ethics Committees of all participating centers. Blood collection, processing, and cfDNA and gDNA extractions were performed as described in a previous publication^12^. Libraries for targeted sequencing were prepared starting from 10-25 ng of cfDNA and 100 ng of sheared gDNA with KAPA HyperPrep Kit (Roche) following the SeqCap EZ HyperCap v2.3 protocol (for hybridization with PCF_SELECT panel v. 2.0) or KAPA HyperCap v3.0 protocol (for hybridization with PCF_SELECT v. 3.0) with a few modifications as per PCF_SELECT panel design^12^. Sample sequencing was performed at the University of Trento by the Next Generation Sequencing Facility on the Illumina NovaSeq 6000 platform with 1000x and 200x intended coverage, respectively, for cfDNA and gDNA.

#### 1kGP WGS (30x)

Sample collection and sequencing protocols are described in the original publication^14^. Samples data included in this study were selected randomly among all super-population groups and sex status to ensure a balanced representation of ethnicities and sex. Fastq files were downloaded from SRA EBI FTP (ftp.sra.ebi.ac.uk/vol1/fastq/) with accession codes provided in **Supplementary Table 1**.

#### CCLE WGS (30x)

Sample collection and sequencing protocols are described in the original publication^15^. Selected cell lines include: 22RV1 and DU145 (prostate cancer), AU565 and BT20 (breast cancer), EF021 and EF027 (ovarian cancer) and CAPAN1 and DANG (pancreatic cancer). Copy number segments were downloaded from CCLE website and coordinates with abs(Segment_Mean) < 0.5 were retained as wild-type segments. Fastq files were downloaded from SRA with accession codes provided in **Supplementary Table 1**.

#### 1kGP long-read WGS (30x)

Sample collection and sequencing protocols are described in the original publication^17^. The list of sample IDs is provided in **Supplementary Table 1**. CRAM files were downloaded from 1KG_ONT_VIENNA repository.

### 5.2 Sequencing data preprocessing

T2T-CHM13 v2.0 reference genome and chain files were downloaded from the UCSC website.

#### Patients targeted (1000x)

Adapters were removed from paired-end reads with trimmomatic^30^ (version 0.32). Reads were aligned with bwa mem (version 0.7.17-r1188) to T2T-CHM13v2.0, G1Kv37 and hg38 FASTA files. SAM files were merged with Picard (version 2.23.9) MergeSamFiles command and converted to BAM format, sorted and indexed with samtools^24^ (version 1.7). BAM realignment and recalibration were performed using GATK^31^ (version 3.8.0). Since GATK resources are not yet available for T2T-CHM13v2.0 reference genome, liftOver of hapmap_3.3.b37.vcf, 1000G_omni2.5.b37.vcf and 1000G_phase1.snps.high_confidence.b37.vcf was performed as explained in Methods 5.3. BAM cleanup was performed with samtools calmd (version 1.7) and overlapping read pairs were clipped through bamUtil (version 1.0.14). SNP pileup and allelic fraction computation were run through PaCBAM^32^ tool (version 1.6.0), providing as input the set of all high MAF SNPs (MAF > 0.2) covered by PCF_SELECT targeted panel^12^. Mapping statistics were obtained with samtools (version 1.7).

#### Breast cancer WES (96-500x), 1kGP WGS (30x) and CCLE WGS (30x)

Adapters were removed from paired-end reads with trimmomatic (version 0.32). Reads were aligned with bwa mem (version 0.7.17-r1188) to T2T-CHM13v2.0 and hg38 FASTA files. SAM files were merged with Picard (version 2.23.9) MergeSamFiles command and converted to BAM format, sorted and indexed with samtools (version 1.7). BAM recalibration was performed using GATK (version 3.8.0) with previously lifted over GATK resources. SNP pileup and allelic fraction computation were run through PaCBAM tool (version 1.6.0), providing as input a set of 30 million high MAF SNPs (MAF > 0.2) interspersed across chromosomes. Mapping statistics were obtained with samtools (version 1.7).

#### 1kGP long-read WGS (30x)

Since CRAM files aligned to hg38 and T2T-CHM13 were downloaded instead of fastq files, no preprocessing was applied. SNP pileup and allelic fraction computation were not performed. Mapping statistics were obtained with samtools (version 1.7) and cramino^33^ (version 1.1.0). Since samtools average quality estimation is not suitable for long-reads, read average quality was computed as the average from the histogram of Phred scale accuracies (corresponding to base qualities) from cramino output.

### 5.3 LiftOver of BED and VCF files

LiftOver of VCF files was achieved through Picard (version 2.23.9) LiftoverVcf function with RECOVER_SWAPPED_REF_ALT = T. LiftOver of BED files was instead performed with UCSC liftOver binary file.

### 5.4 VCF files preprocessing and VAF computation

Raw genotype calls for all 1kGP 30x samples were downloaded for hg38 and T2T-CHM13v2.0 as VCF files (description on how the files were produced is provided in the original publications^2,14^) from [1kGP FTP website](https://ftp.1000genomes.ebi.ac.uk/vol1/ftp/data_collections/1000G_2504_high_coverage/) and [T2T consortium repository](https://s3-us-west-2.amazonaws.com/human-pangenomics/T2T/CHM13/assemblies/variants/1000_Genomes_Project/chm13v2.0/all_samples_3202/). Variant allele frequency (VAF) was computed from genotype calls allelic depth (AD) field as following: alt_count / (alt_count + ref_count), where ref_count and alt_count correspond to AD first and second field, respectively. VCF files were filtered for retaining only entries with at least 10 reads of coverage support and heterozygous VAF (0.2 ≤ VAF ≤ 0.8). Only SNPs included in the set of 30 million high MAF SNPs interspersed across chromosomes (used in Methods 5.2 for SNP pileup) were kept for RMB analyses.

### 5.5 Theoretical VAF distribution

Theoretical VAF distributions were obtained by simulating VAF values at multiple coverage bins ([5, 20], [20,30], [30,50], [50,100], [100,200]). Random sampling of allele A and allele B from a set of balanced alleles was performed for each coverage bin, maintaining a consistent per-bin SNP numerosity with respect to the hg38 patients’ WES data.

### 5.6 Reference mapping bias analysis

VAF curves were produced by selecting all heterozygous SNPs (0.2 ≤ VAF ≤ 0.8) with at least 20 supporting reads. For CCLE WGS pileups, only SNPs overlapping wild-type segments were considered. Distribution peaks were established by selecting the closest peak to 0.5 computed through localMaxima function from MsCoreUtils package. Allele swapping classification was attributed when the reference allele of hg38 assembly was equal to the alternative allele in T2T-CHM13 assembly and viceversa. SNPs with heterozygous VAF in T2T-CHM13 or hg38 correspond to calls where only one of the assemblies passes both the coverage and heterozygosity threshold. Delta VAF was computed as | *T*2*T VAF* − *hg*38 *VAF* | and SNP classification was performed with respect to 0.005 threshold (delta VAF < 0.005 and delta VAF ≥ 0.005). VAF difference among T2T-CHM13 and hg38 was tested with a one-tailed paired t-test with parameter alternative = “greater” comparing T2T-CHM13 VAF distribution with hg38 VAF distribution of SNPs with delta VAF > 0.005, referred throughout the text as “VAF incoherent”.

### 5.7 Read mapping analysis

Coordinates for a set of 3085 SNPs with concordant classification across at least 3 WES samples were converted to BED format (both for hg38 and T2T-CHM13 reference genome) and used to subset each WES whole blood sample’s BAM file through bedtools bamtobed and intersect functions (version v2.25.0). Output QNAMEs were then given as input through READ_LIST_FILE field to Picard (version 2.23.9) FilterSamReads function to extract data related to read mapping for reads of interest from original BAM files aligned to hg38 and T2T-CHM13.

SNP support was assessed based on bedtools intersect file, which matches each SNP rsID with a set of QNAMEs. Only reads mapping to a unique SNP were filtered to compare mapping features. Read classification was performed according to chromosome mapping (either multiple mapping, different chromosome, same chromosome, or uniquely mapped to either hg38 or T2T-CHM13). Multiple mapping reads were excluded from the analysis. SNP concordant support status was determined when QNAMEs assigned to the SNP were equal in hg38 and T2T-CHM13, hence SNPs with discordant support have a differential support of at least one read.

### 5.8 SNP proximity sequence analysis

The analysis of each SNP’s proximity was performed by converting its coordinates (chr and position) to a bed format specified as chromosome, position – 150 and position + 150 for hg19, hg38 and T2T-CHM13 reference genomes. The FASTA sequence of each interval was obtained from Biostrings getSeq function^34^ giving as input the BS reference genome file (BSgenome.Hsapiens.NCBI.T2T.CHM13v2.0, BSgenome.Hsapiens.UCSC.hg19 and BSgenome.Hsapiens.UCSC.hg38) and assembly-specific genomic coordinates. The set of random SNPs was selected through createRandomRegions function of gUtils R package with input BSgenome.Hsapiens.NCBI.T2T.CHM13v2.0 and a GenomeRanges including all SNP coordinates, telomeres and centromeres to mask. Output ranges were lifted over to hg38 through liftOver function using hs1 (T2T-CHM13) to hg38 chain file.

Similarity of DNA sequences was assessed by pairwise comparison of FASTA sequences across assemblies (T2T-CHM13 versus hg19, T2T-CHM13 versus hg38 and hg19 versus hg38) through the pwalign R package^35^ through the function compareSequences().

GC content was computed as the percentage of G and C bases within the analyzed sequence as:

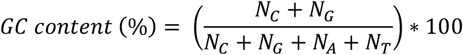

Repetitiveness of DNA sequences was measured through a custom R implementation of the D_2_^R^ algorithm^36^. The output value represents a proxy of repetitiveness, where low values correspond to low k-mer repetition (k-mer size = 3). Since the threshold for repetitiveness detection must be defined heuristically, values were compared directly without imposing a pre-determined threshold.

### 5.9 FASTA sequence extraction and pairwise sequence alignment

FASTA sequences were obtained with twoBitToFa UCSC utility^29^ providing as input a specific bed file for each track and the.2bit file for the reference genome of choice. Latest version. 2bit files were downloaded from UCSC golden path repositories for hg19, hg38 and hs1 reference genome (GRCh37.p13, GRCh38.p14 and T2T-CHM13v2.0 version, respectively). Pairwise sequence alignment was performed through seq-align Needleman and Wunsch algorithm implementation (https://github.com/noporpoise/seq-align) and formatted with a custom script to store the alignment result as a compact strings. Specifically, each pairwise alignment is summarized as a pipe delimited string where each event is expressed as a symbol, the event starting position and parentheses holding the number of bases undergoing the event. The symbol “?” corresponds to mismatches, “+” to insertions and “-” to deletion, assigned according to the first of the two sequences. For example, the following alignment is summarized as ?626(1)|-627(1)|+630(2):

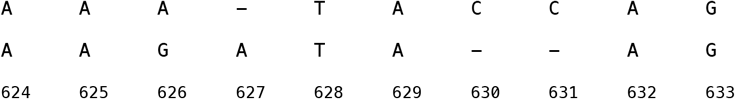

In addition to the summarized string, the custom script keeps track of the alignment score produced by Needleman and Wunsch algorithm and the total number of mismatches, insertions and deletions.

### 5.10 CpG islands annotation and methylation analysis

T2T-CHM13 v2.0 UCSC CpG unmasked track was downloaded from UCSC data repository. T2T-CHM13 CpG islands bed regions were lifted over to hg19 and hg38 (Methods 5.3), obtaining a set of common N = 49388 CpG islands.

CpG sequence analysis was performed as presented in Methods 5.9. In addition, GC content (computed as in Methods 5.8) and the position of each C base computed for each CpG and compared among assemblies to obtain a granular classification highlighting CpGs where the number of Cs or their position varies.

Reduced representation bisulphite sequencing (RRBS) fastq files from 5 CCLE cancer cell lines (PANC0504, DU4475, OAW28, MDAPCA2B and HS606T) were analyzed through Nextflow methylseq^37^ pipeline (version 2.6.0), both with T2T-CHM13 and hg38 as FASTA reference genomes. Bismark methylation calls for CpG context were converted to read tables through a custom script. Resulting tables were then filtered to retain only common reads and at least 4 methylation sites in hg38-aligned table.

### 5.11 ENCODE TFBS peaks annotation and TF enrichment analysis

hg38 ENCODE Transcription Factor ChIP-seq Clustered track (encRegTfbsClustered) was downloaded from UCSC data repository. encRegTfbsClusteredWithCells UCSC track for hg38 was lifted over to T2T-CHM13 and hg19 (Methods 5.3). TFBS peaks sequence analysis for hg19, hg38 and T2T-CHM13 was performed as presented in Methods 5.9.

The TFBS track was then filtered to retain only entries with support in at least 5 different cell lines, obtaining a set of 8 transcription factors (MYC, SP1, MAFK, CEPBP, FOXA1, FOS, JUND and CTCF). Motif enrichment analysis was performed through ame_rank() function from MEME suite R implementation^38^ by giving as input the whole set of sequences supporting peaks for a specific gene and the reference genome (hg19, hg38 and T2T-CHM13). The enrichment rank output was then stored for each transcription factor and compared among assemblies.

### 5.12 ClinVar variant annotation

dbSNP version 155 and ClinVar version 20220313 VCF files were downloaded from NCBI archive for GRCh37 and GRCh38 assembly. LiftOver of dbSNPv155 and ClinVar version 20220313 VCF from GRCh38 to CHM13v2.0 were downloaded from [T2T-CHM13 GitHub repository](https://github.com/marbl/CHM13).

ClinVar VCF files were indexed and divided by chromosome, retaining only entries mapping to chromosomes 1:22, X and Y. Chromosomes in RefSeq accession format in dbSNP VCFs were converted through NCBI sequence report files. For comparing differences across assemblies, chromosome VCF files were merged by unique IDs, maintaining only entries present in all three VCF files.

As per SNP proximal sequence analysis, each variant’s coordinates (chr and position) were converted to a bed format specified as chromosome, position – 150 and position + 150 for hg19, hg38 and T2T-CHM13 reference genomes. Results are provided in Supplementary Table 4.

Pairwise alignment outcome was formatted and assigned to a classification based on the presence or absence of mismatches according to different thresholds:

- Allele swapping: the reference base and alternative base of the variant under study are opposite among the two reference genomes;
- [1,5] mismatches: among 1 and 5 mismatches;
- > 5 mismatches | indels: more than 5 mismatches or presence of insertions and/or deletions (indels);

Actionability classification was performed by intersecting gene names with genes included in COSMIC Actionability v14^22^ signature.

### 5.13 Variant calling over ClinVar sites

Variant calling was performed on all tumor samples (N=28, with N=15 cfDNA prostate cancer samples sequenced with 3Mb panel and N=13 cfDNA breast cancer samples sequenced with WES) providing as input a VCF with all ClinVar sites for hg38 and T2T-CHM13 reference genome, respectively. Variant callers used for the analysis are state-of-the-art methods Strelka2 and BCFtools. Strelka2 was run with default parameters with and without Manta indel calling and –indelCandidates parameter (referred to as Strelka2.indel and Strelka2.noindel, respectively). BCFtools was run with mpileup command, specifying-d 2000 and-Q 20, and call command with-A-m parameters. Comparison of outcome calls was performed by integrating all VCF outputs based on their position and sample of origin. Variant allele frequency (VAF) was computed from genotype calls allelic depth (AD) field as following: alt_count / (alt_count + ref_count), where ref_count and alt_count correspond to AD first and second field, respectively.

### 5.14 BLAST analysis of gene nucleotide sequences

FASTA sequences of genes of interest were obtained through twoBitToFa UCSC utility^29^ providing as input a specific bed file for each track and the.2bit file for the reference genome of choice. The nucleotide sequence of each gene was compared with blastn command line tool with T2T-CHM13 as query and hg38 as subject. The output was then parsed and visualized through seqvisr^39^ R package. Exon coordinates for hg38 were downloaded from UCSC Table Browser^29^ GENCODE v47 knownGenes track.

### 5.15 IGV-style visualization of sequencing reads

BAM files were extracted with Rsamtools^40^ function and filtered to divide reads supporting the alternative allele from all other reads spanning the region of interest (±150 bp from the variant). Coordinates for the chromosome bands were downloaded from goldenPath website for each reference genome. Figures were built using Gviz^41^ R package and BSgenome^42^ libraries, with custom modification of Gviz functions.

### 5.16 Comparison of read consensus with reference sequence in long-read 1kGP samples

SNVs with differences in the proximal sequence among hg38 and T2T-CHM13 from Supplementary Table 8 were filtered for pathogenic and likely pathogenic variants and support of VAF or coverage change in at least one sample, resulting in a total of N = 76 SNVs. Read consensus was obtained with samtools consensus (version 1.22) command by giving as input a bed with the chromosome, SNV coordinate – 150 and SNV coordinate + 150 separately for hg38 and T2T-CHM13 aligned samples and with-q parameter equal to 20. The consensus sequence for each SNV in each sample was then directly compared to the reference genome of choice with a custom R script. Specifically, Levenshtein similarity was computed through comparator^43^ R package among the FASTA sequence consensus and FASTA sequence extracted from BSgenome^42^ libraries with the same bed coordinates.

## 6 Resources availability

### 6.1 Lead contact

Requests for further information and resources should be directed to and will be fulfilled by the lead contact, Francesca Demichelis.

### 6.2 Data and code availability

- Fastq, CRAM, and VCF files for cell lines and 1kGP samples were downloaded from publicly available repositories (Methods 5.1).
- BAM files of 3Mb panel sequencing (prostate cancer) and WES (breast cancer) samples are available upon request.
- Annotated pairwise alignment tables for ClinVar, CpG islands and TFBS are provided on a Zenodo repository (10.5281/zenodo.17953686).
- OpenCRAVAT annotation for ClinVar can be found among [OpenCravat Modules](https://run.opencravat.org/submit/nocache/index.html?store) under the name “Clinvar T2T-hg38 comparator annotator”.
- The code and material for reproducing the figures is available on GitHub (https://github.com/demichelislab/ref_gen_comp).
- A set of scripts used for sequence analysis of UCSC tracks and homopolymers is available on GitHub (https://github.com/demichelislab/annotating_change_ref_genomes).

## 7 Author contributions

I.C., F.O., Y.C., and F.D. conceptualized and designed the study. Y.C. and F.D. supervised the study’s overall conduct. I.C. performed computational data analysis and statistical analyses. O.Q. prepared sequencing libraries. M.P. contributed materials for data analysis. F.D. was responsible for the funding acquisition for this study. I.C., F.O., Y.C., and F.D. wrote the original draft of the manuscript. I.C., F.O., O.Q., M.P., Y.C., and F.D. reviewed and edited the manuscript.

## 9 Acknowledgments

We thank the CIBIO Department Next-Generation Sequencing (NGS) Facility. We also thank all authors and consortia who contributed to the public datasets we queried, including the Cancer Cell Line Encyclopedia (CCLE), the T2T consortium, and the 1000 Genome Project. Funding support was provided by the Accelerator Award 2018, funded by Cancer Research UK (A26822) and Fondazione AIRC per la ricerca sul cancro ETS (22792), Fondazione Cassa Di Risparmio Di Trento E Rovereto, and the Department of CIBIO – University of Trento. Y.C. is supported by PCF Challenge Award 2024 to F.D.

