## Supplementary Figures for "T2T-CHM13 reference genome reduces mapping bias and enhances alignment accuracy at disease-associated variants"

### List of Supplementary Figures

Figure S1. Alignment to T2T-CHM13v2.0 reference genome improves several mapping metrics across diverse sequencing designs

Figure S2. Reference Mapping Bias analysis of short-read sequencing data

Figure S3. Comparison of mapping features from reads spanning a set heterozygous SNPs in WES gDNA samples

Figure S4. Detected variation within CpG islands and transcription factor binding sites sequences has a negligible impact on methylation calling and transcription factor (TF) enrichment analysis

Figure S5. Genomic proximities of ClinVar variants differ among reference genomes

Figure S6. Detection of clinically relevant SNVs with annotated sequence discrepancies is enhanced in samples aligned to T2T-CHM13

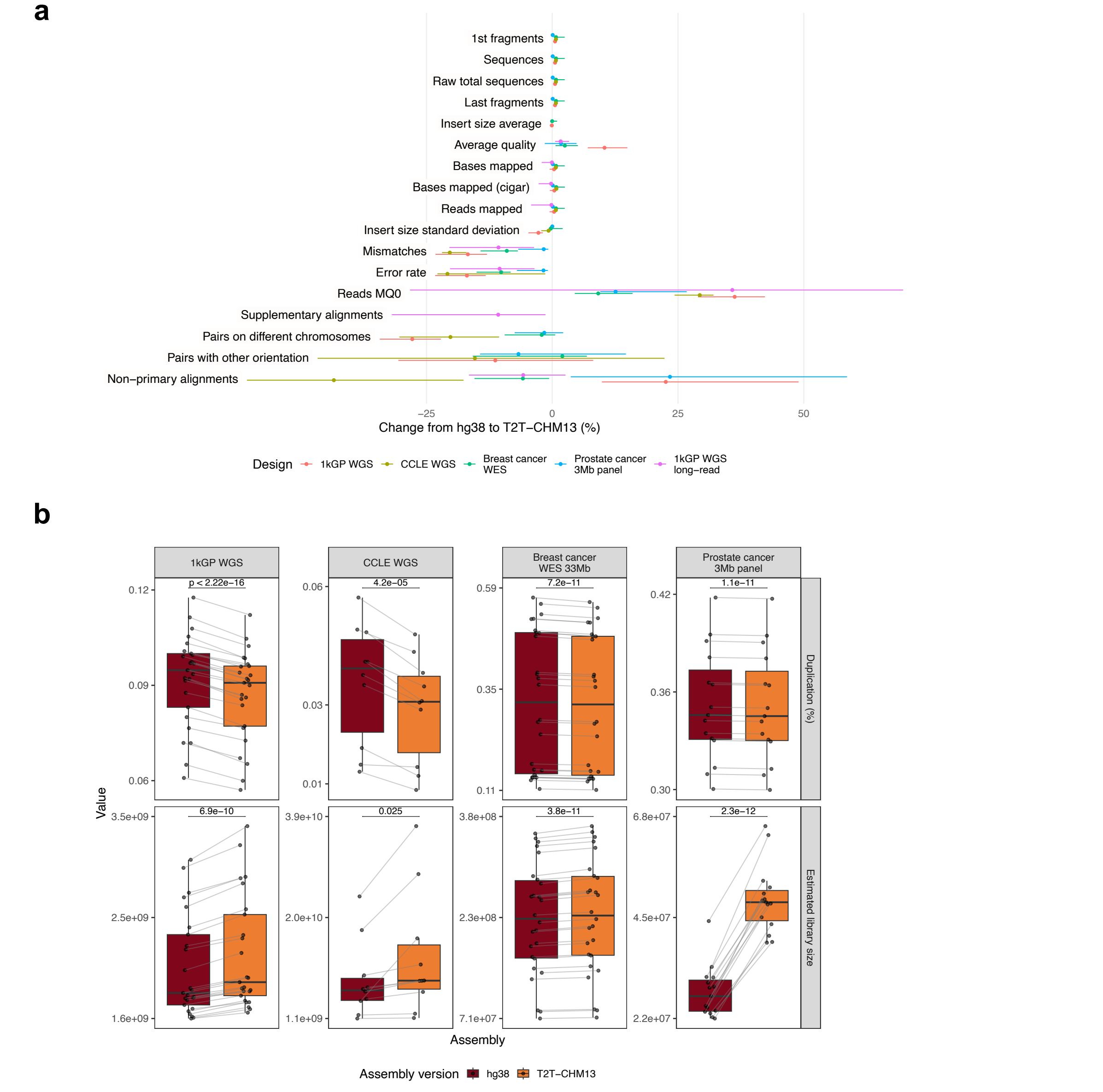


**Supplementary Figure 1 | Alignment to T2T-CHM13v2.0 reference genome improves mapping metrics across diverse sequencing designs.** a) Summary of samtools stats metrics for all analyzed samples (N = 1106). Median (dot) and min-max (line) values of relative change from hg38 to T2T-CHM13 are colored according to sequencing design. b) Paired Picard statistics for samples aligned to hg38 and T2T-CHM13 for multiple short-read sequencing designs. Paired t-test p-values were computed for each design and metric combination. Data points from the same sample are connected by gray lines.

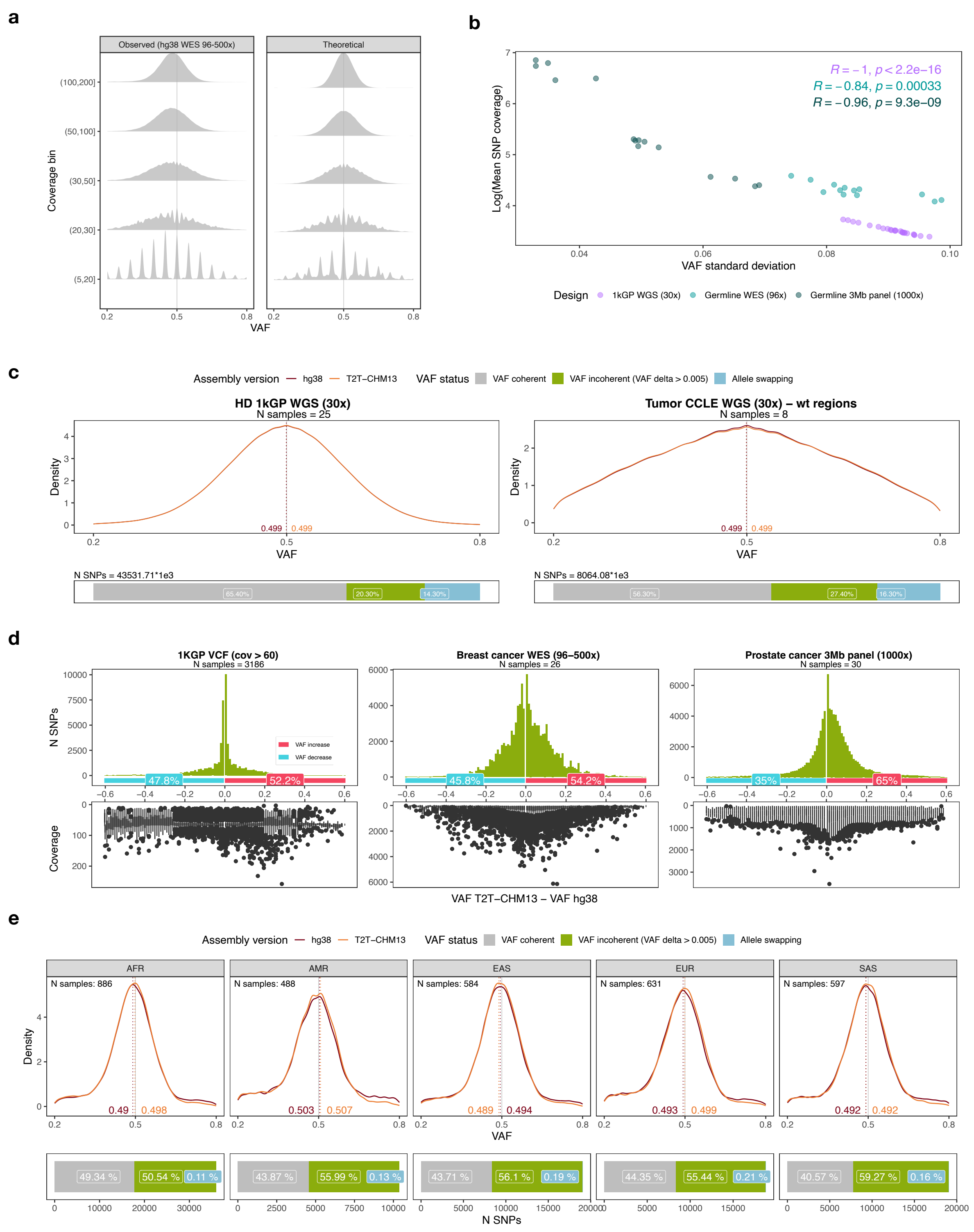


**Supplementary Figure 2 | Reference mapping bias (RMB) analysis of short-read sequencing data.** a) Distribution of observed hg38 VAF from patients WES data and theoretical VAF at multiple coverage bins (Methods 5.5). The continuous gray vertical line at

0.5 VAF indicates the expected heterozygous value. b) Correlation among VAF standard deviation and mean coverage (log scale) over heterozygous SNPs for germline samples included in short-read 1kGP WGS, WES, and targeted 3Mb panel sequencing data. Pearson correlation R value and p-value reported for each design. c) Reference mapping bias analysis for 1kGP WGS (left) and CCLE WGS (right) samples. The top panel includes all heterozygous SNPs VAF values and reports the numeric value of each density peak, corresponding to the dotted vertical lines. The underlying barplot provides a classification for each SNP based on VAF comparison among hg38 and T2T-CHM13. d) Mirrored histogram and boxplots report information on SNPs with delta inconsistent VAF identified from 1kGP VCF analysis. The histogram includes an annotation with the percentage of entries with VAF increase (pink) and VAF decrease (blue). e) VAF peaks (top) and barplot (bottom) of 1kGP VCF heterozygous SNPs with at least 60 supporting reads divided by super-population groups (AFR = African, AMR = Ad Mixed American, EAS = East Asian, EUR = European, SAS = South Asia). Top: densities of heterozygous SNPs VAF values with numeric value of each density peak, corresponding to the dotted vertical lines. The expected VAF value of 0.5 is indicated by the continuous gray vertical line. The number of analyzed samples per superpopulation is indicated in each facet. Bottom: barplot providing the classification for each SNP based on VAF comparison among hg38 and T2T.

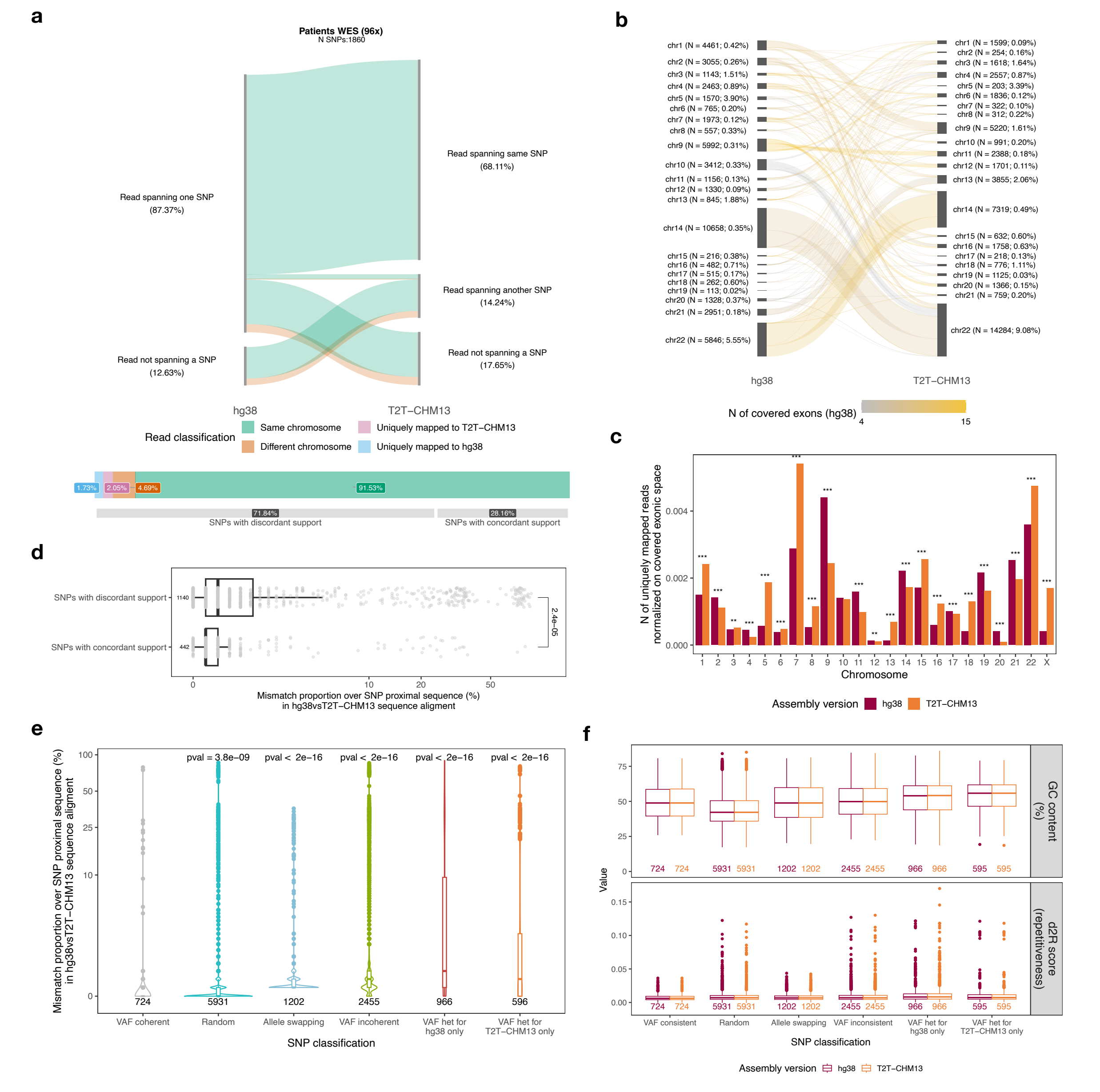


**Supplementary Figure 3 | Comparison of mapping features from reads spanning a set heterozygous SNPs in WES gDNA samples.** a) Sankey plot for read classification among T2T-CHM13 and hg38. The color coding indicates mapping features of each read. Column breaks highlight the SNP support of each read in both assemblies. The annotations on the bottom summarize the percentage of reads by read mapping classification and the percentage of SNPs with concordant and discordant read support. b) Sankey plot depicting reads that change chromosome mapping among hg38 and T2T-CHM13 alignment. For each chromosome, the number of reads and the percentage of exome sequence they cover with respect to the whole chromosome's exonic space is reported in parentheses. Flows are colored according to the number of exons covered by the reads. c) Per-chromosome count of uniquely mapped reads to T2T-CHM13 and hg38 assembly, normalized on each chromosome's exonic space. Proportion test significance is reported for each chromosome (* = p-value < 0.05; ** = p-value < 0.01; *** = p-value < 0.001; **** = p-value < 0.0001). d) Mismatch proportion distribution computed on the proximal sequence of each SNP (± 150 bp). The number of observations included in each boxplot and one tailed Wilcoxon test p-value are reported on the left and right of the plot, respectively. X-axis values are pseudo-log transformed. e) Proportion of mismatches detected in SNP proximity (± 150 bp) hg38vsT2T-CHM13 sequence comparison. Distributions for each SNP classification category encompass a subset of SNPs analyzed in RMB comparison analysis (Supplementary Table 3). “Random” group includes a set of randomly selected genomic sequences (301 bp length). Wilcoxon test p-value was computed with respect to consistent VAF group. f) Direct comparison of GC content and d2R repetitiveness score for each SNP classification category. Numerosity is reported for each category. Wilcoxon paired test was performed for each paired boxplot without reaching statistical significance.

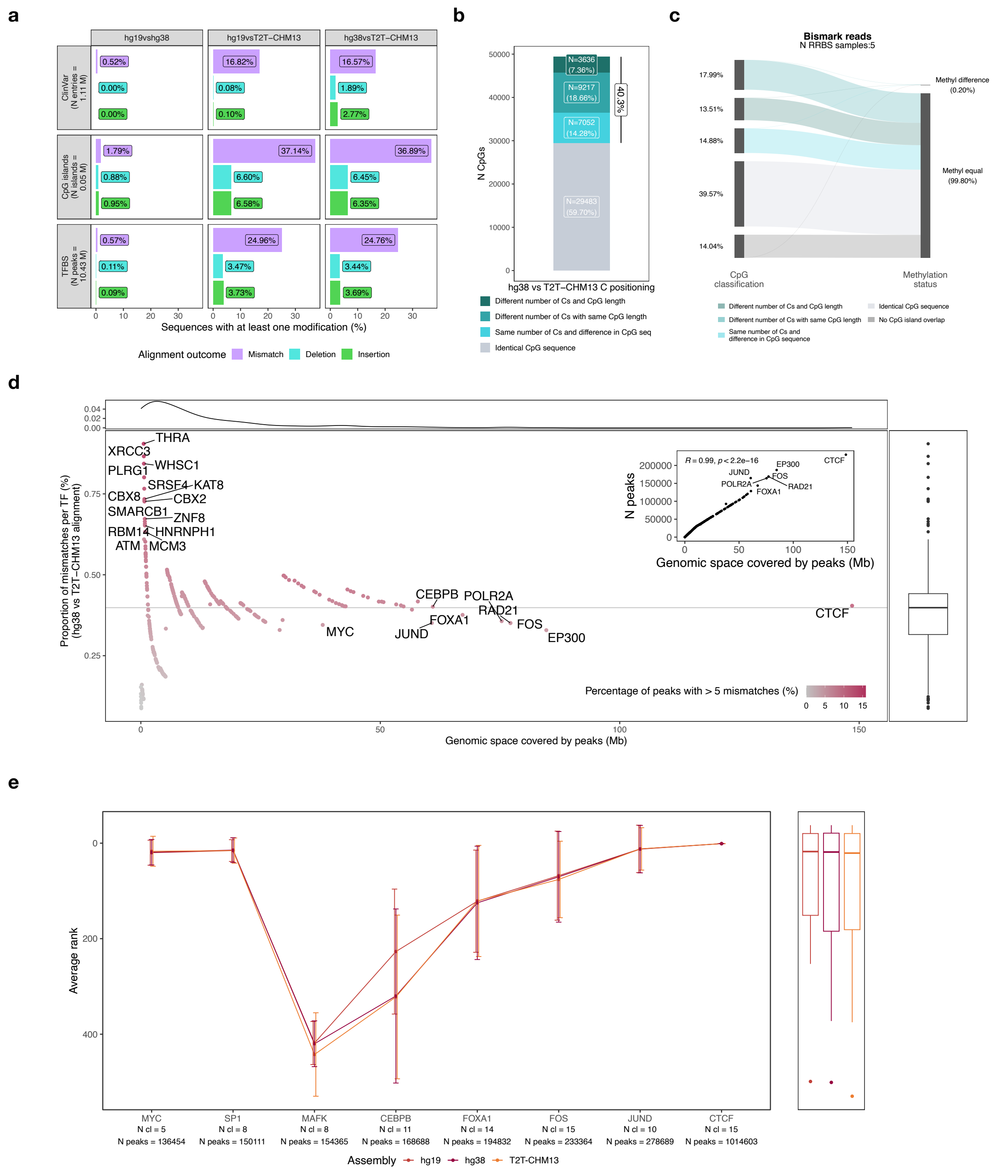


**Supplementary Figure 4 | Detected variation within CpG islands and transcription factor binding sites sequences has a negligible impact on methylation calling and transcription factor (TF) enrichment analysis.** a) Barplot displaying the proportion of track entries with at least one modification (insertions, deletions and mismatches) in pairwise sequence alignment among hg19, hg38 and T2T-CHM13. b) CpG island classification according to hg38 versus T2T-CHM13 sequence alignment outcome. Each bar reports the number of CpGs and the percentage of each category with respect to the overall analyzed set. c) Sankey plot of Bismark read methylation status of 5 RRBS tumor cell lines among hg38 and T2T-CHM13 with respect to CpG classification. d) Scatter plot reporting the proportion of mismatches for hg38vsT2T-CHM13 alignment compared to the genomic space covered by the peaks of each transcription factor. Each labelled dot is colored according to the proportion of peaks with > 5 mismatches over the total number of peaks per TF, with a darker shade corresponding to a higher proportion. Dark gray continuous horizontal line highlights the median proportion value across TFs. Summary distributions are reported for x- and y-axis. The inset plot shows the number of analyzed peaks per TF and genomic space covered by peaks with Pearson correlation coefficient and p-value annotation. e) Motif enrichment analysis (AME) for transcription factors supported in at least 5 cell lines. Ranks for the intended TF were extracted from TF enrichment analysis outcomes and are reported as error bars (average ± standard deviation). Bottom labels on x-axis state the number of supporting cell lines (N cl) and the number of peak sequences (N peaks) used in AME enrichment per TF. Summary boxplots for the y-axis compare the average rank among assemblies.

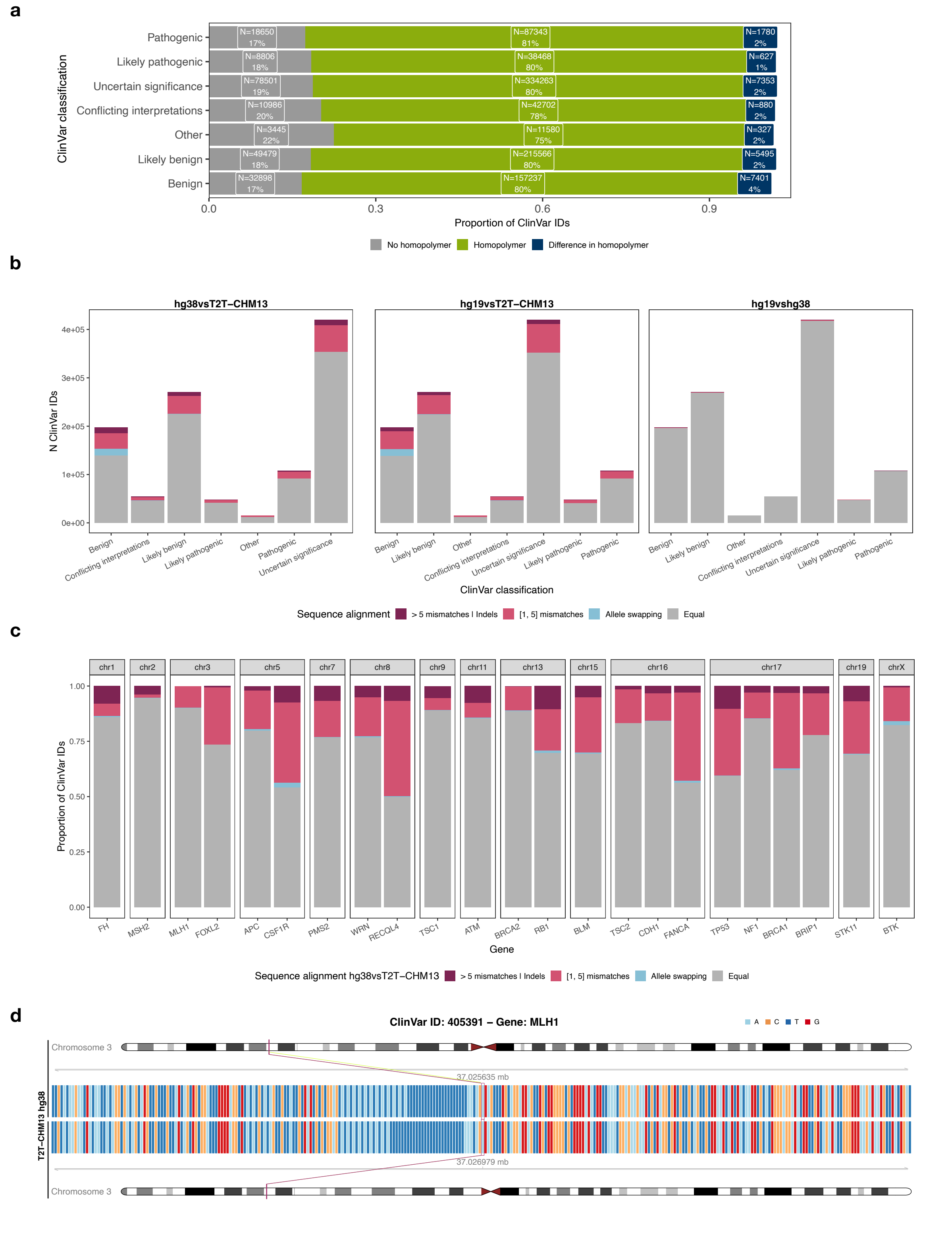


**Supplementary Figure 5 | Genomic proximities of ClinVar variants differ among reference genomes.** a) Proportion of ClinVar entries with homopolymers (at least 5 consecutive bases) in their proximal sequence. b) ClinVar IDs stratification based on pairwise sequence alignment classification for hg38 versus T2T-CHM13, hg19 versus T2T-CHM13 and hg19 versus hg38 comparison. c) Subset of actionable genes with at least 5% of pathogenic variants with a modified sequence among hg38 and T2T-CHM13, divided by chromosome and sorted by genomic position (hg38 coordinates). d) Example of neighboring sequence variation among hg38 and T2T-CHM13 for a ClinVar site over MLH1 gene. The SNV is highlighted in red and its genomic position is reported both numerically in Mb and on the chromosome ideogram for both assemblies.

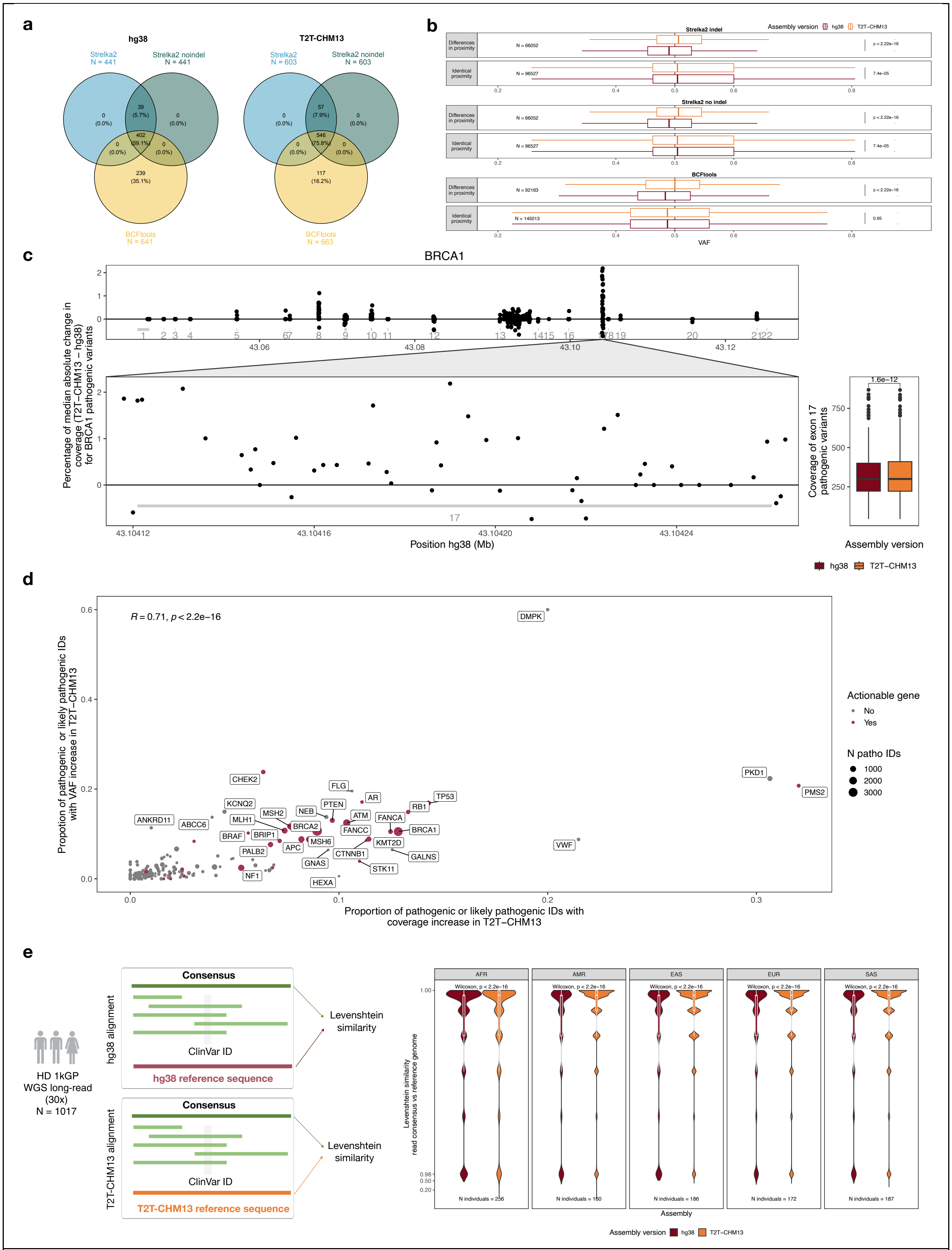


**Supplementary Figure 6 | Detection of clinically relevant SNVs with annotated sequence discrepancies is enhanced in samples aligned to T2T-CHM13.** a) Intersection among SNV calling outputs for Strelka2 with InDel correction, Strelka2 without InDel correction and BCFtools. Calls were defined as ClinVar annotated SNVs with VAF >= 5%, coverage > 50 and quality > 200. b) Boxplot for VAF estimation from variants with and without variation in their 150 bp proximity for each variant caller. Wilcoxon test p-value was computed for all comparisons. Numerosity is reported for each boxplot. c) Median delta coverage (coverage T2T-CHM13 – coverage hg38) among all analyzed samples across *BRCA1* gene pathogenic variants. Exon number and segments are depicted in dark gray. The zoom on *BRCA1* exon 17 highlights an increase in coverage at most pathogenic variants in the region for T2T-aligned samples. Boxplots on the right report coverage values for each analyzed sample and p-value of one-tailed paired Wilcoxon test for T2T-CHM13 > hg38. d) Proportion of pathogenic and likely pathogenic variants with coverage and/or VAF increase by gene. Dot sizes are proportional to the number of pathogenic and likely pathogenic variants present within each gene. e) Read consensus analysis over pathogenic SNV proximal sequences in N=1017 long read samples from 1kGP project. Left: sketch of the read consensus analysis. Right: distribution of Levenshtein similarity computed among read consensus per patient over pathogenic SNVs proximal sequences and the reference genome sequence to which samples were aligned by 1KGP super-population groups (AFR = African, AMR = Ad Mixed American, EAS = East Asian, EUR = European, SAS = South Asia). Y-axis is log transformed for values below 0.9 and expanded non-linearly for values greater than 0.9. One-tailed paired Wilcoxon test for hg38 < T2T p-values and the number of samples per super-population are reported on the top and bottom of each facet, respectively.
